# Slow breathing impacts inter-organ dynamics modulating brain function and risk behavior

**DOI:** 10.64898/2026.01.09.698695

**Authors:** Wenhao Huang, Mine Schmidt, Ignacio Rebollo, Felix Molter, Min Pu, Beatrix Keweloh, Lok Yan Lam, Peter N.C. Mohr, Gabriele Bellucci, Stefan Röttger, Soyoung Q Park

## Abstract

Successful decision-making requires that external information be interpreted in the context of the body’s state. Within the framework of body-brain interaction, deliberately modifying one’s autonomic state can shape how we evaluate the world, ultimately influencing choices. Yet, it remains unclear whether and how intentional autonomic regulation affects human decision-making. In this study, we tested instructed prolonged exhalation, a slow-breathing technique designed to boost parasympathetic activity during risky decision-making. Participants followed distinct breathing protocols while making risky choices, with neural and physiological activity measured using fMRI and multi-channel monitoring. Prolonged exhalation increased risky choices by enhancing reward sensitivity and elevating cardiac parasympathetic activity. Importantly, individuals with greater parasympathetic upregulation also showed stronger reward-related responses in the ventromedial prefrontal cortex and precuneus. Our work reveals a transformative role for breathing-based interventions, demonstrating that breathing-based autonomic regulation can shape value-based decision-making through neuro-cardiac pathways.

**GRAPHICAL ABSTRACT:** 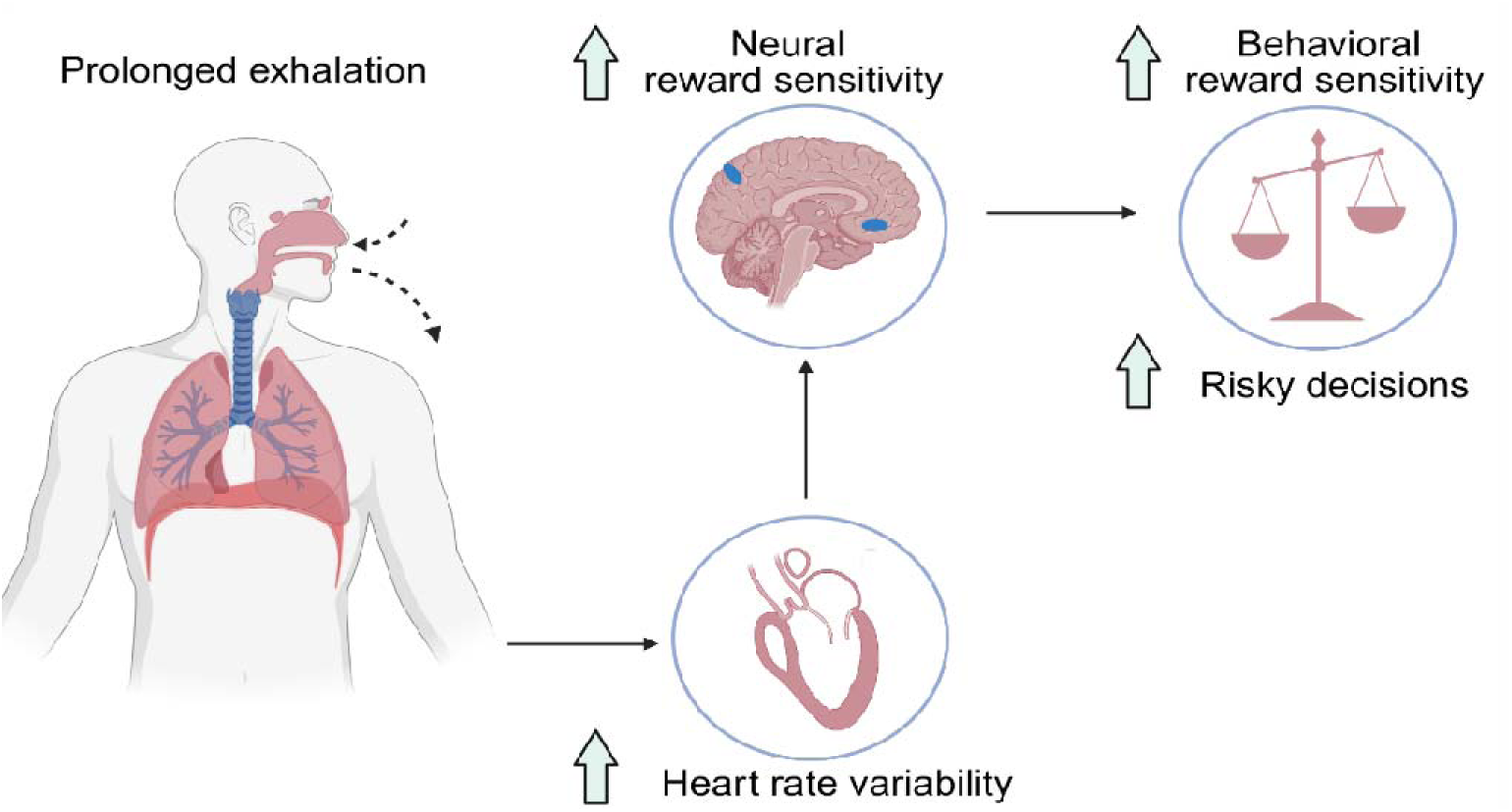

**HIGHLIGHTS:**

- Prolonged exhalation increases risky decisions by increasing the reward sensitivity
- Prolonged exhalation enhances cardiac parasympathetic activity without dampening sympathetic activities
- Greater cardiac parasympathetic activity under prolonged exhalation amplifies neural reward sensitivity in the ventromedial prefrontal cortex and precuneus

## INTRODUCTION

Imagine you’re running late and rushing to the bank. Upon arrival, you’re immediately faced with an important investment decision. Here, your momentarily unrelated physiological arousal state, such a increased heart rate or rapid breathing, influences your choice, potentially leading to suboptimal decision you might later regret. However, the same choice made calmly, under a more relaxed physiological state, may lead to a satisfying result. This raises the crucial question: Can we regulate our own physiological responses to make better decisions?

Prolonged exhalation is a breathing technique, with a shorter inhalation to longer exhalation ratio (e.g., 2:8 seconds), that has been shown to effectively shift the cardiac parasympathetic predominance^1,2,3^. Thi is driven by the tight synchronization of breathing with cardiac timing, such that during inhalation, the heart rate increases, whereas during exhalation it decreases, resulting in periodic fluctuations in heart rate, called heart rate variability (HRV)^4,5^. A prolonged exhalation phase enhances these fluctuations and thus leads to an overall increase in HRV, reflecting respiration-coupled cardiac vagal modulation^6,7,8,9^.

This breathing technique becomes particularly crucial in the framework of intentional decision enhancement. A typical decision option consists of both rewards and losses under uncertainty, with which we guide our choices^10,11,12^. Importantly, rewards and losses are interpreted differently, depending on the body’s current internal autonomic state^13,14^: States of cardiac parasympathetic predominance (often described as rest-and-digest) have been associated with greater sensitivity to potential rewards, wherea sympathetic predominance (fight-or-flight) has been linked to relatively greater sensitivity to potential losses. Supporting this, individual differences in cardiac parasympathetic activity, reflected by higher resting HRV, are associated with greater reward sensitivity. In the Balloon Analogue Risk Task, individuals with higher HRV(high-frequency HRV (HF-HRV)) are more inclined to take reward-related risk^15^. Similarly, people with greater resting HF-HRV have a stronger preference for advantageous decks in the Iowa Gambling Task, indicative of increased sensitivity to long-term rewards^16^. In contrast, enhanced sympathetic activity, as reflected in pupil dilation and skin conductance, has been associated with heightened sensitivity to loss and increased risk-avoidant behavior^17,18^. However, it is unknown whether a voluntary upregulation of cardiac parasympathetic activity via regulated breathing enhances momentary risky decisions or reward sensitivity.

At the neural level, the ventromedial prefrontal cortex (vmPFC) is a strong candidate linking cardiac parasympathetic predominance and reward processing. The human vmPFC not only encodes the subjective value of choice options^12,19^ but is also embedded within the central autonomic network^20^, receiving interoceptive input from the anterior insula^21^, and participating in the integration of interoceptive and affective states^22,23^. In line with this, individual differences in HRV (indexed by SDNN (standard deviation of inter-beat intervals), a time-domain measure) were shown to predict changes in vmPFC activity in a task exerting self-control from food reward^24^. Despite this evidence pointing toward prolonged exhalation as a potential modulator of decisions, its feasibility and its exact neuro-visceral mechanism are unknown.

The current study aims to systematically investigate whether and how prolonged exhalation causally impacts decision-making. We hypothesized that prolonged exhalation (1) increases risky choices, (2) enhances cardiac parasympathetic activity, and (3) changes neural reward representation (pre-registered hypotheses https://osf.io/4cbfz).

We employed a within-subject experimental procedure, in which participants (n=41; 24 female, mean ± SD age = 24.78 ± 4.93 years) performed a risky choice task while breathing either according to prolonged exhalation or eupnea (natural respiration pattern) in a counterbalanced manner. Both breathing patterns were instructed visually by dynamically increasing and decreasing bars, during which participants made decisions^11^ (Figure 1). In each trial, participants were asked to either accept or reject a gamble. Each gamble had a 50% probability of depicted rewards and losses (Figure 1C). During the entire session, participants’ brain activity was measured by means of functional magnetic resonance imaging (fMRI). At the same time, we assessed physiological markers of parasympathetic and sympathetic activities (i.e., respiration, cardiac activity, skin conductance, and pupil responses).

**Figure 1.**
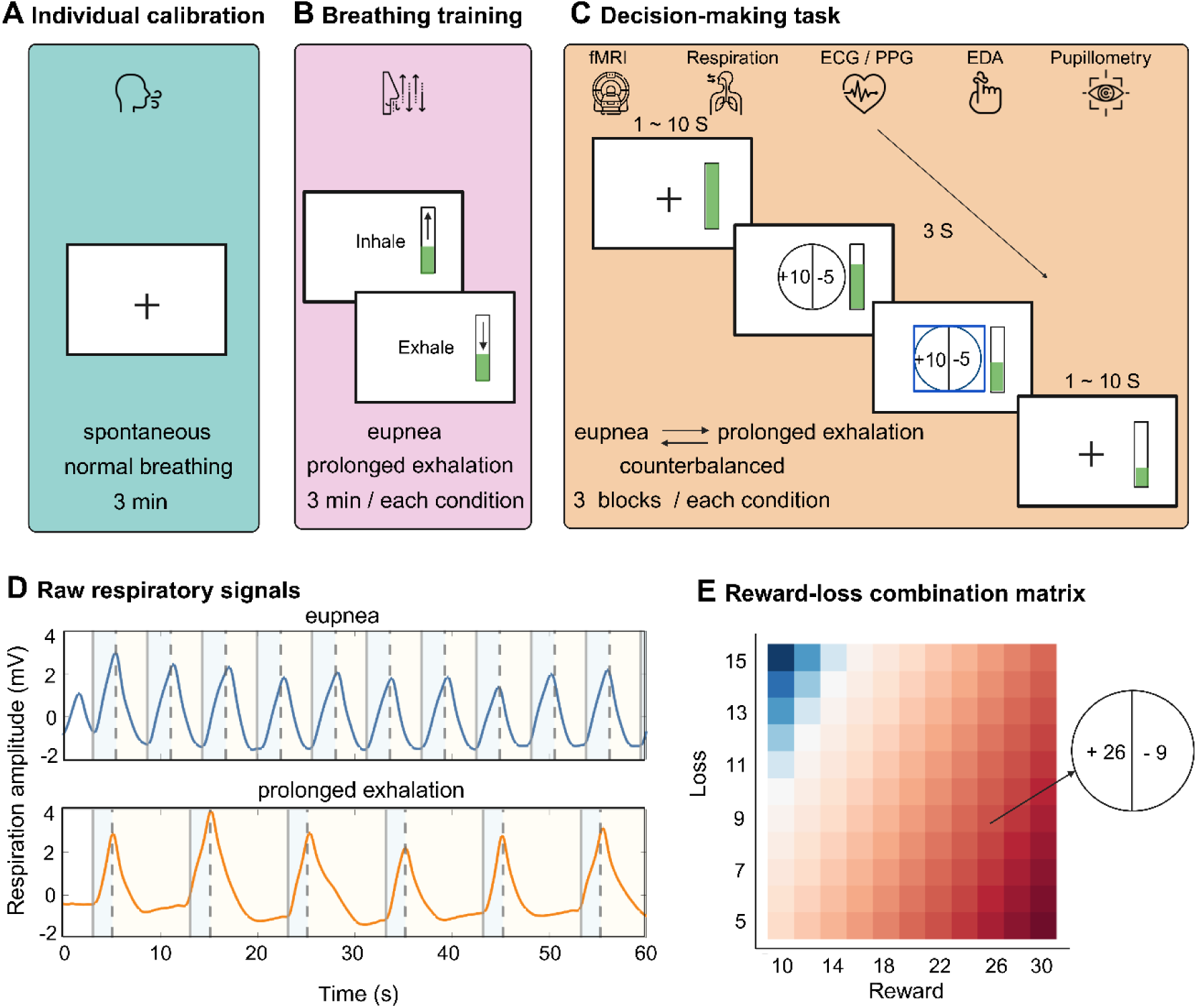
Experimental procedure. (A) First, individual natural breathing pattern was assessed for individualized eupnea breathing instruction. (B) Participants practiced instructed breathing for both prolonged exhalation and eupnea breathing, presented by visual cues. Both breathing patterns were instructed, to keep the cognitive load consistent. (C) Participants either accepted or rejected an option consisting of a certain amount of reward and loss with a 50% probability of realization while following the instructed breathing pattern displayed as a progress bar on the right. Breathing conditions were presented in three-block sets, and the order of these sets was randomly counterbalanced across participants. (D) Raw respiratory data of the two breathing conditions eupnea (blue) and prolonged exhalation (orange) of one exemplary participant. Vertical grey solid lines depict the instructed inhalation onset. Vertical dashed lines depict the instructed exhalation onset. During eupnea, participants followed their natural breathing rhythm, while during prolonged exhalation, the ratio was fixed at 2:8 seconds. (E) For each trial, the reward and loss values were systematically selected from a predefined reward-loss matrix. Rewards ranged from €10 to €30 (in steps of €2), and losses from €5 to €15 (in steps of €1), 120 unique combinations used across both conditions. Red and blue shading indicate positive and negative expected values of a gamble (EV = 0.5 × reward − 0.5 × loss), respectively, and white indicates gambles with an expected value close to zero.

## RESULTS

### Validation of breathing manipulation

We first confirmed that participants followed the breathing instructions by testing whether the breathing conditions modulated exhalation duration and respiration rate on average using paired t-tests (see Method). Exhalation duration was significantly longer in the prolonged exhalation condition compared to eupnea (t(40) = 34.621, p < 0.001, Cohen’s d = 5.407; mean difference = 5.183 s, SE = 0.150, 95% confidence intervals, CI = [4.880, 5.485]; Figure 2A). In line with this, the respiration rate was significantly lower in the prolonged exhalation condition compared to eupnea (t(40) = -11.616, p < 0.001, Cohen’s d = - 1.814; mean difference = -6.104 breaths/min, SE = 0.525, 95% CI = [-7.166, -5.042]; Figure 2B). These results confirm that the prolonged exhalation manipulation successfully slowed the respiratory rhythm by extending the duration of exhalation.

**Figure 2.**
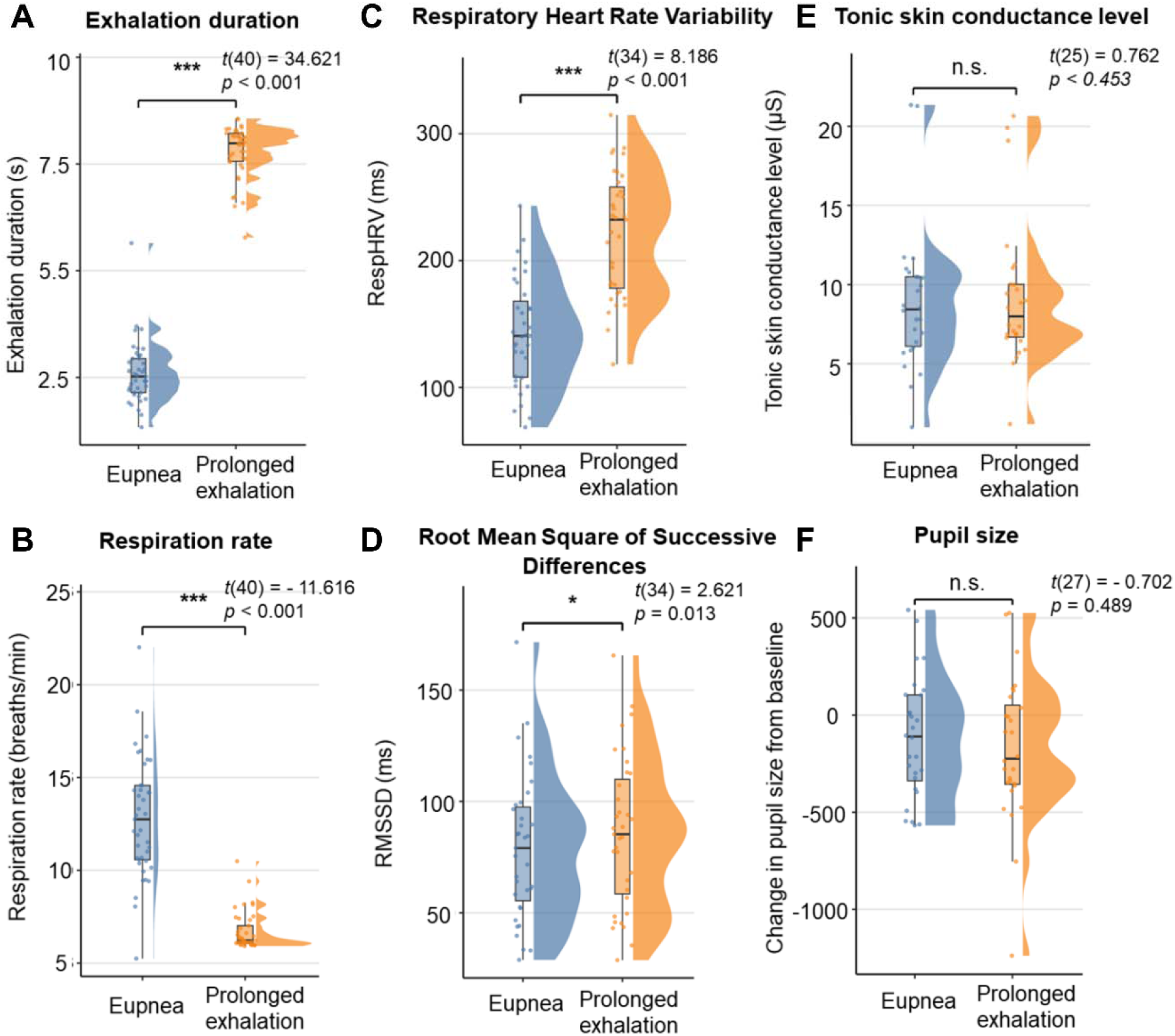
Effects of prolonged exhalation on physiological markers of parasympathetic and sympathetic activity. Prolonged exhalation (A) significantly increased exhalation duration and (B) significantly decreased respiration rate, compared to eupnea. Also, cardiac parasympathetic activity was enhanced as (C) Respiratory Heart Rate Variability (RespHRV) and (D) Root Mean Square of Successive Differences (RMSSD) of heart rate variability were significantly higher during prolonged exhalation compared to eupnea. In contrast, sympathetic read-outs, such as (E) tonic skin conductance level, and (F) change in pupil size from baseline showed no significant difference between conditions, indicating prolonged exhalation selectively modulates cardiac parasympathetic activity. * *p* < 0.05, *** *p* < 0.001; n.s., not significant.

### Prolonged exhalation enhances cardiac parasympathetic activity

We next tested whether prolonged exhalation enhances parasympathetic activity, by two well-established cardiac indicators^25,26^, Respiratory Heart Rate Variability (RespHRV)^27^ also known as Respiratory Sinus Arrhythmia (RSA) and Root Mean Square of Successive Differences (RMSSD). RespHRV is a direct index of cardiorespiratory coupling^8,28^, whereas RMSSD is a time-domain index of cardiac parasympathetic modulation, which quantifies rapid beat-to-beat variability in heart rate. It is less sensitive to task-related breathing variability, optimal for a stable cardiac estimation in experimental contexts^26,29^.

Prolonged exhalation significantly enhanced cardiac parasympathetic activity, reflected by higher RespHRV (t(34) = 8.186, p < 0.001, Cohen’s d = 1.384; mean difference = 78.000 ms, SE = 9.529, 95% CI = [58.636, 97.364]; Figure 2C) and RMSSD (t(34) = 2.621, p = 0.013, Cohen’s d = 0.443; mean difference = 7.012 ms, SE = 2.675, 95% CI = [1.575, 12.449]; Figure 2D) compared to eupnea.

We also examined sympathetic indices, namely tonic skin conductance level and pupil size changes. Interestingly, neither tonic skin conductance level (t(25) = 0.762, p = 0.453, Cohen’s d = 0.149; mean difference = 0.273 µS, SE = 0.358, 95% CI = [-0.465, 1.011]; Figure 2E), nor pupil size (t(27) = - 0.702, p = 0.489, Cohen’s d = - 0.133; mean difference = - 57.016, SE = 81.212, 95% CI = [-223.649, 109.617]; Figure 2F) differed significantly between conditions. These results suggest that the prolonged exhalation selectively increased cardiac parasympathetic activity without inducing a general shift in sympathetic arousal.

### Prolonged exhalation increases risky choices and selectively enhances reward sensitivity

We next examined whether prolonged exhalation modulates risky choices, by a generalized linear mixed-effects model (GLMM) with a logit link function to predict trialwise binary decisions (accept = 1 vs. reject = 0) from breathing condition (prolonged exhalation = 1, eupnea = -1), reward, loss of the trial, and their interactions with breathing condition. The model included subject-level random intercepts and random slopes for breathing condition, reward, and loss. Notably, prolonged exhalation significantly increased trial-wise risky choices, as observed by the main effect of breathing condition on trial-wise choices (β = 0.168, SE = 0.071, z = 2.37, p = 0.018, 95% CI = [0.0292, 0.306]; Figure 3A). Importantly, this increase in risky choices was driven by reward sensitivity, as under prolonged exhalation, reward had a greater impact on the decisions, compared to eupnea. This was unveiled by the significant breathing condition × reward interaction (β = 0.176, SE = 0.053, z = 3.32, p < 0.001, 95% CI = [0.0719, 0.279]; Figure 3A). Post hoc comparison confirmed that the reward slope reflecting how strongly reward magnitude predicted decisions was significantly steeper under prolonged exhalation than under eupnea (β = 0.351, SE = 0.106, z = 3.32, p < 0.001; Figure 3B), suggesting enhanced reward sensitivity under prolonged exhalation.

**Figure 3.**
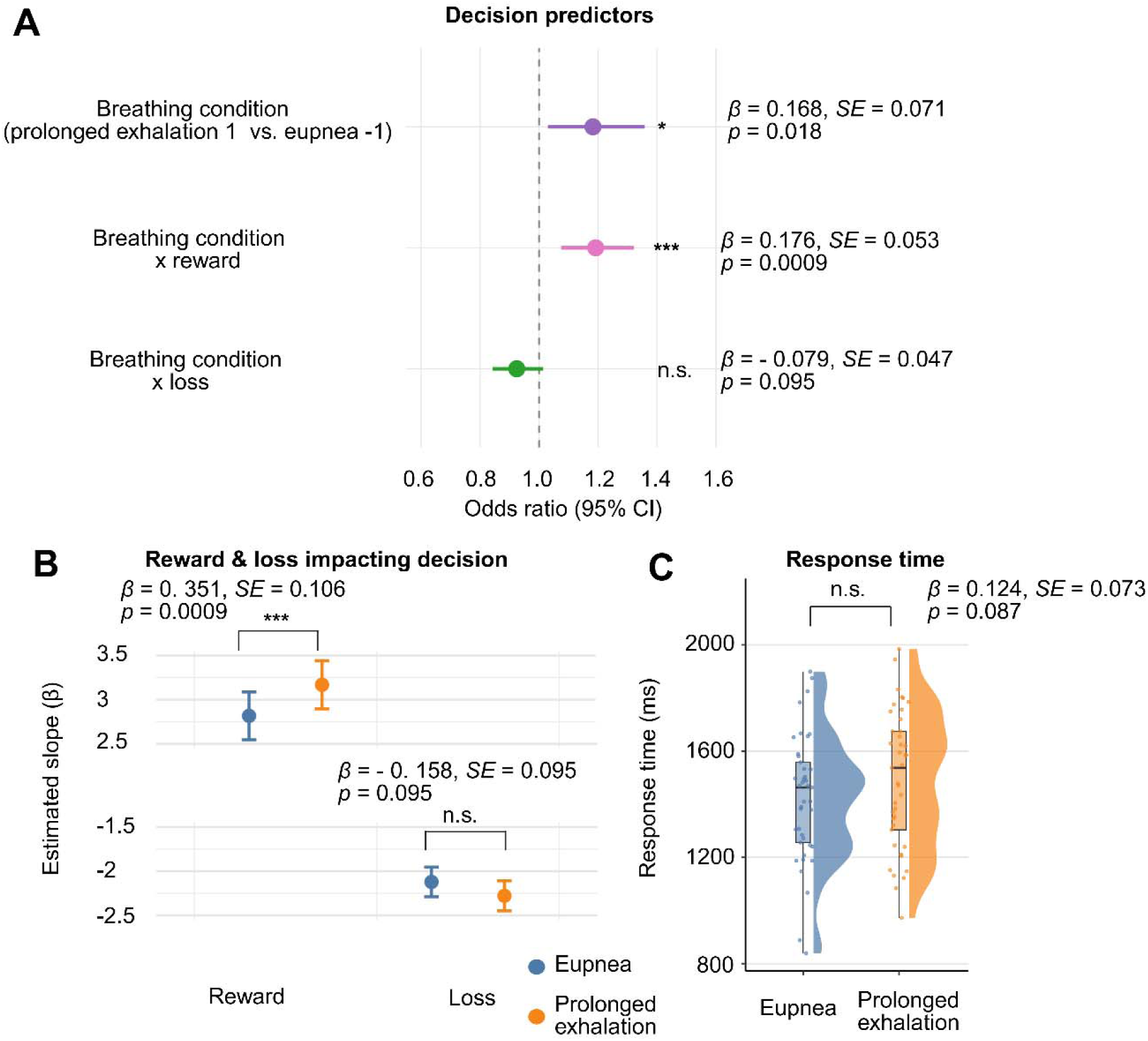
Prolonged exhalation increases the impact of reward on decisions. (A) Under prolonged exhalation participants generally increased risky decisions significantly more often and the impact of reward on decisions increased, observed by a significant condition × reward interaction. Points depict odds ratios with 95% confidence intervals. The dashed line indicates no effect (OR = 1). (B) Reward impact on decision was significantly greater under prolonged exhalation, compared to eupnea, as model-estimated reward slopes were significantly higher under prolonged exhalation than eupnea. Error bars represent ±1 SE of model-estimated slopes. (C) Response times did not differ significantly between breathing conditions. Points represent participant-level mean RTs; densities and boxplots depict group-level distributions and interquartile ranges. * *p* < 0.05, *** *p* < 0.001; n.s., not significant.

Critically, the impact of prolonged exhalation was specific to the influence of reward on decisions, as we observed no change in the influence of loss on decisions, as the condition × loss interaction did not reach significance (β = -0.079, SE = 0.047, z = -1.67, p = 0.095, 95% CI = [-0.171, 0.0138]; Figure 3A, B).

To directly test whether prolonged exhalation differentially modulated reward versus loss sensitivity, we formally contrasted the condition × reward and condition × loss interaction terms using a likelihood-ratio test. This comparison revealed a significant asymmetry (χ²(1) = 7.73, p = 0.005, Table S6), indicating that prolonged exhalation more strongly modulated sensitivity to reward than to loss. We additionally examined whether prolonged exhalation modulated individual loss aversion by estimating the prospect-theory loss aversion parameter (λ). No significant difference in λ was observed between breathing conditions (Figure S3A, B)^10,11^. Importantly, a parameter-recovery analysis (Table S10) indicated limited identifiability of the inverse temperature parameter (recovery r = 0.23) in this accept–reject design. Given this constraint, parameter-level estimates should be interpreted with caution. Accordingly, we treat the prospect-theory parameters as descriptive and base our main inferences on the trial-wise GLMM analyses, which directly test value sensitivity without relying on the estimation of trade-off–prone latent parameters. Response time did not differ significantly between breathing conditions (β = 0.124, SE = 0.073, z = 1.71, p = 0.087; Figure 3C). Importantly, supplementary analyses indicate that these decision-making patterns are not attributable to reduced decision noise, inattention, or response-time–mediated effects (Tables S1–S4).

### Respiration-driven cardiac parasympathetic shifts amplify neural reward encoding

We next investigated whether and how this cardiac parasympathetic shift under prolonged exhalation reshapes the neural representation of reward during decisions. To do so, we specified a general linear model (GLM) that predicted blood-oxygen-level-dependent (BOLD) response with parametric modulation of trial-wise reward and losses of decision options. Then, at the group level, individual first-level contrast images reflecting reward-related brain activation were entered into a whole-brain regression analysis, with individual differences in breathing condition-dependent HRV (prolonged exhalation vs. eupnea) as a continuous predictor (RMSSD). We found that individuals showing larger cardiac shifts (ΔRMSSD) under prolonged exhalation also exhibited stronger reward-related activity in the vmPFC (Figure 4A & Figure 4C; MNI coordinates: x = -6, y = 39, z = -18; t(33) = 4.79, thresholded at cluster-level p < 0.05, family-wise error (FWE)-corrected) and the precuneus (Figure 4B & Figure 4D; MNI coordinates: x=0, y=−51, z=60; t(33) = 4.23, FWE_c_ < 0.05) during choices. All fMRI analyses were conducted after removing the respiratory and cardiac variance from the data to rule out the possibility that the observed effects are driven by the general physiological changes (see Method).

**Figure 4.**
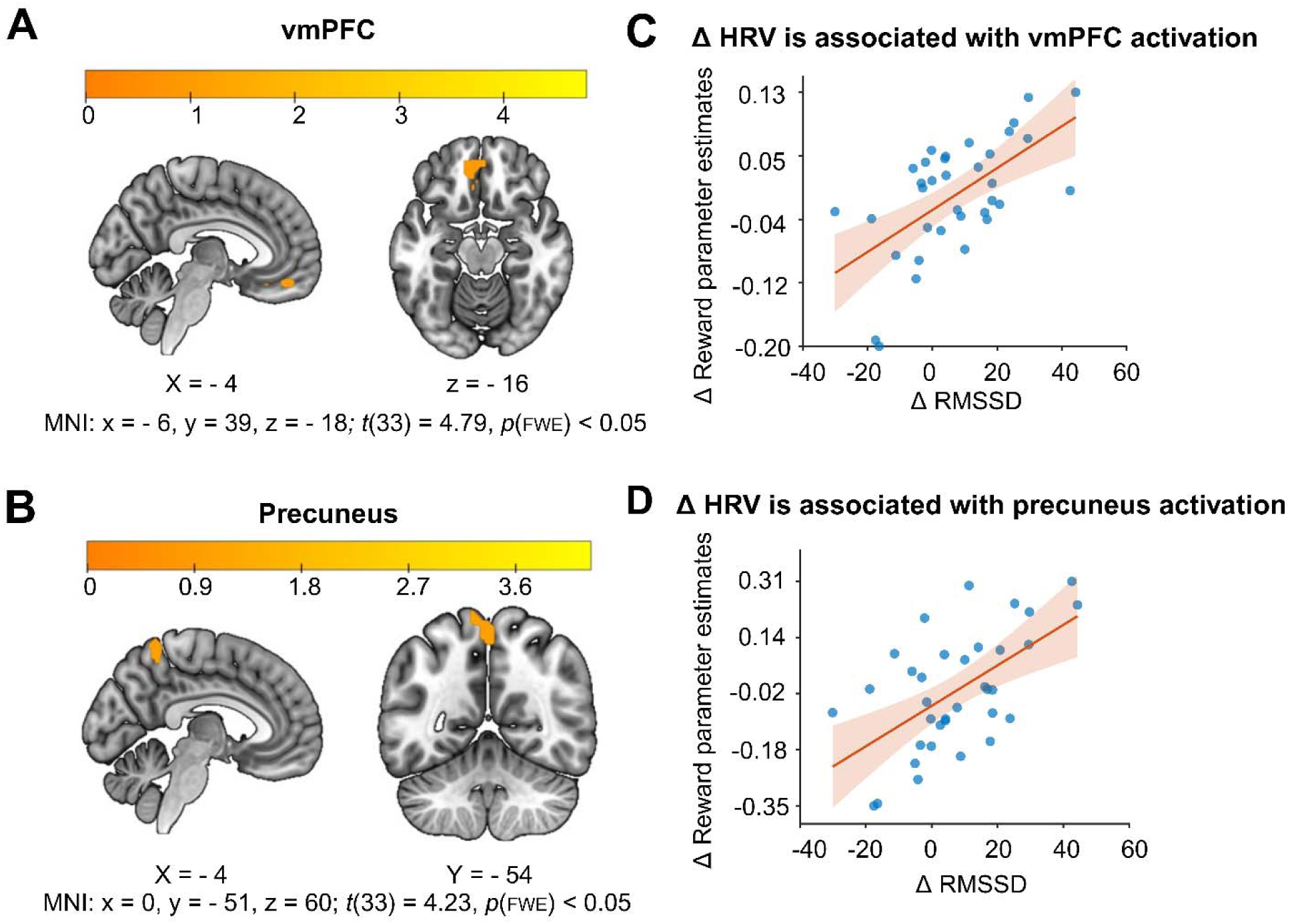
Parasympathetic modulation of reward-related neural activity under prolonged exhalation. Individuals showing larger cardiac parasympathetic shifts under prolonged exhalation also exhibited stronger reward-related activity in the (A) vmPFC and (B) precuneus. Our whole-brain regression analysis revealed a significantly positive association between changes in cardiac parasympathetic activity (ΔRMSSD = RMSSD (prolonged exhalation - eupnea)) and reward-related activation in these regions. Scatter plots (C, D) show individual relationships between ΔRMSSD and reward-related parameter estimates extracted from the vmPFC and precuneus, respectively.

## DISCUSSION

Here, we provide evidence for a neuro-visceral pathway through which prolonged exhalation i associated with systematic changes in autonomic state, value-related brain activity, and risky choice. To do so, we applied a multimodal study, integrating instructed breathing with simultaneous multi-channel physiological recordings and functional neuroimaging during decision-making. Our task independentl varied reward and loss magnitudes with a 2:1 ratio to yield balanced choice distributions, providing sufficient variability to detect breathing-related effects on value-based decision-making. We demonstrate that prolonged exhalation significantly increases cardiac parasympathetic markers and selectively enhances rewards’ impact on decisions without altering sensitivity to losses. We also observed a significant main effect of prolonged exhalation, which increased the proportion of risky decisions. Strikingly, fMRI analyses revealed that on the group level, individual differences in breathing-induced cardiac parasympathetic enhancement was significantly associated with reward-related BOLD activation in the vmPFC and precuneus at the individual level. Our multimodal approach provides integrated insights into the neurophysiological behavioral framework to understand how bodily changes can result in the modulation of the brain’s reward processing, resulting in enhanced risky choices.

While previous research focused on individual differences in ANS variability^15,16^, we demonstrate how changes in the breathing pattern acutely impact parasympathetic activity and thereby neural reward sensitivity during decision-making. Our results not only extend the existing literature on body-brain interaction^1,24^ but also open a new avenue for exploring how breathing can be used to voluntarily control autonomic states and directly influence decision-making processes. Our investigation uniquely contributes to the current decision neuroscience literature, which mainly focuses on the external situation changes leading to decision modulation. Instead, these data suggest that external stimuli, such as reward, are interpreted in the light of momentary internal bodily state, in this case cardiac vagal tone, which is a crucial factor that needs to be considered.

In our study, we employed a prolonged exhalation breathing technique. Most existing slow breathing protocols focus primarily on maintaining a constant slow respiratory rate (e.g., ∼6 breaths/min) rather than explicitly emphasizing extended exhalation phases. In contrast, prolonged exhalation emphasizes a longer exhalation-to-inhalation ratio, which is physiologically meaningful because baroreflex-mediated cardiac vagal activation occurs primarily during exhalation when temporary increases in arterial pressure trigger baroreceptor activity^30,31^. Importantly, compared to other slow breathing techniques, prolonged exhalation is particularly easy to learn because there is no need to fully standardize the breathing rate, allowing participants to breathe at their own pace. This contributes to restoring cognitive function while benefiting from the parasympathetic effects of prolonged exhalation. Confirming this, Röttger and colleagues have shown that the parasympathetic-enhancing effect of prolonged exhalation does not trade off cognitive performance^3^. In direct comparison, other breathing techniques (i.e., tactical breathing) reduced cognitive performance. In line with this, our prolonged exhalation breathing protocol heightens parasympathetic indices without a downregulation of sympathetic markers (e.g., skin conductance, pupil diameter), suggesting a selective or partial modulation of autonomic branches^8,29^, which potentially also contributes to the preserved cognitive function and decision-making. This aspect is particularly important in the framework of decision-making, in which the primary goal is to make effective choices and not mere relaxation.

Contrasting the traditional decision-theoretic view, we show that the evaluation of potential rewards and losses depends on bodily arousal states^32,33^. Specifically, heightened loss sensitivity and more cautious decision-making have been associated with increased sympathetic activation, as reflected in physiological markers such as heart rate and skin conductance during stress or threat exposure^33,34^. Conversely, cardiac parasympathetic states, such as higher baseline HRV or enhanced cardiac vagal activity, have been shown to be associated with more balanced reward-loss evaluation, better emotional regulation, and stronger cognitive control^35,36,37^. Empirical findings also highlight that cardiac parasympathetic predominance can foster adaptive decision processes: individuals with elevated HRV show reduced susceptibility to the framing effect^38^. Our data build upon these insights and provide a neuro-visceral account, by demonstrating an actively increased cardiac vagal tone via prolonged exhalation not only promotes a calmer physiological state but also enhances the weight assigned to potential rewards during choice. Importantly, this shift does not compromise decision quality; instead, it is in line with prior proposals that autonomic regulation may support more differentiated reward evaluation^15,16^.

At the neural level, individual differences in cardiac parasympathetic enhancement (indexed by ΔRMSSD) predicted reward-related BOLD activation in two key cortical regions: the vmPFC and the precuneus. The vmPFC is well-established as a core hub for subjective-value representation and motivational integration^19,39^. By contrast, the precuneus is implicated in self-referential thought^36,40^ and acts as a bridge between internal bodily signals and one’s mental simulation^41^. Its integrative role suggests a potential link between physiological states and self-relevant evaluation, consistent with evidence that individual differences in precuneus connectivity with vmPFC or dlPFC are associated with how people value immediate versus delayed rewards^42^. Together, these activations suggest a targeted neural mechanism in which prolonged exhalation driven parasympathetic upregulation aligns with enhanced reward valuation in the vmPFC. The precuneus may integrate bodily signals into the higher-order cognitive processes, such as mental simulation and self-relevant evaluation, that shape value-based decision-making. Our exploratory analysis showed that questionnaire-based interoceptive self-regulation scores were associated with task-based precuneus activity specifically (Figure S4), but not with vmPFC activation, further supporting its relation in bodily self-integration. Interestingly, our analyses did not reveal correlations between parasympathetic enhancement and the insula or anterior cingulate cortex (ACC), regions commonly associated with interoception^13,43^. However, unlike these studies showing a general link between interoception and activation in the insula and ACC, our analysis specifically focuses on the impact of a respiration-modulated increase in cardiac parasympathetic activity on risky decisions, rather than interoception in general.

Although prior work has implicated striatum/midbrain and insula/amygdala in reward and loss processing, we did not observe robust breathing-related modulation in these regions. Instead, breathing-related effects were confined to cortical regions involved in subjective value representation, most prominently the vmPFC^12^. We believe this pattern reflects two features of the present design. First, our analyses focused on value computation during the option phase and did not include an outcome phase that typically elicits strong prediction-error or loss-evoked signals in subcortical circuits associated with learning-based value updating^12^. Second, prolonged exhalation primarily induces a parasympathetic state shift, which may preferentially bias cortical value integration processes rather than engage classical subcortical or aversive systems. Consistent with this interpretation, prolonged exhalation did not alter behavioral loss sensitivity, aligning with the absence of breathing-related effects in loss-related regions such as the insula or amygdala^11^.

Taken together, these findings suggest that the observed cortical pattern does not reflect a generalized effect on learning or affective processing, but can instead be interpreted within a broader framework of brain–autonomic integration. In particular, the Neurovisceral Integration Model^22^ proposes that prefrontal regions provide a key interface through which autonomic state can influence higher-order cognitive evaluations. From this perspective, respiration-driven parasympathetic modulation may bias value-related processing by altering the cortical context in which reward information is evaluated, rather than by directly engaging subcortical learning or affective circuits.

Exploratory analyses examining whether individual differences in ProEx-induced autonomic changes (ΔRMSSD, ΔSCL, Δpupil) predicted individual differences in behavioural change (Δacceptance) did not reveal strong or consistent associations (Table S12). This suggests that the breathing-induced parasympathetic shift is more clearly reflected in trial-wise neural reward encoding (vmPFC/precuneus) than in coarse, condition-averaged changes in risky choice. Accordingly, the relationship between autonomic modulation and behaviour may be more strongly expressed at the level of neural value representation, rather than through a direct coupling between tonic autonomic state and risk preference.

The selective impact of prolonged exhalation breathing on reward responsiveness has important implications for clinical contexts, such as anxiety, panic disorder, and depression, given their distinct autonomic signatures and maladaptive reward processing^44,45,46^. By enhancing cardiac parasympathetic modulation through prolonged exhalation techniques, individuals may restore reward processing, a valuable pathway for emotional recalibration^44,47^. Prolonged exhalation harbors the potential for a low-cost, low-risk, easily applicable intervention to be incorporated into therapy or rehabilitation programs, especially to support pharmacological treatments.

Beyond clinical settings, sustaining parasympathetic engagement under acute stress may support value-based decision-making in high-risk professional contexts, such as emergency response, elite sports, military operations, and aviation, where decisions are made under intense pressure and uncertainty. Prolonged exhalation has been shown to reinforce emotional resilience and cognitive performance^3,48^ while enhancing parasympathetic activation. Given prolonged exhalation’s portability and simplicity, it may be particularly well-suited for on-the-spot or field applications where real-time arousal regulation is mandatory. Scientifically, our results have crucial implications for neuroscience, economics, psychology, and psychiatry, fields in which physiological data, such as heart rate and respiration, are often considered as ‘noise’ and neglected. Our findings add to a growing body of work on body–brain and autonomic–decision interactions, and highlight the value of incorporating physiological measures into decision neuroscience.

Taken together, our findings indicate that prolonged exhalation selectively enhances parasympathetic activity and heightens neural reward representation, thereby increasing reward sensitivity and biasing choice toward accepting gambles. This pattern underscores the essential function of body-state in shaping human decisions, an empirical validation for neurovisceral integration^22,35^. Our findings refine current theories of body-brain interaction, by revealing effective and easy-to-implement strategies regulating physiological functions, such as breathing and nutrition^49,50^, to intentionally impact the body and brain to change decisions. Our study suggests potential applications for breathing-based interventions, though future work is needed to test their efficacy across diverse populations and real-world situations.

## STAR⍰METHODS

### KEY RESOURCES TABLE

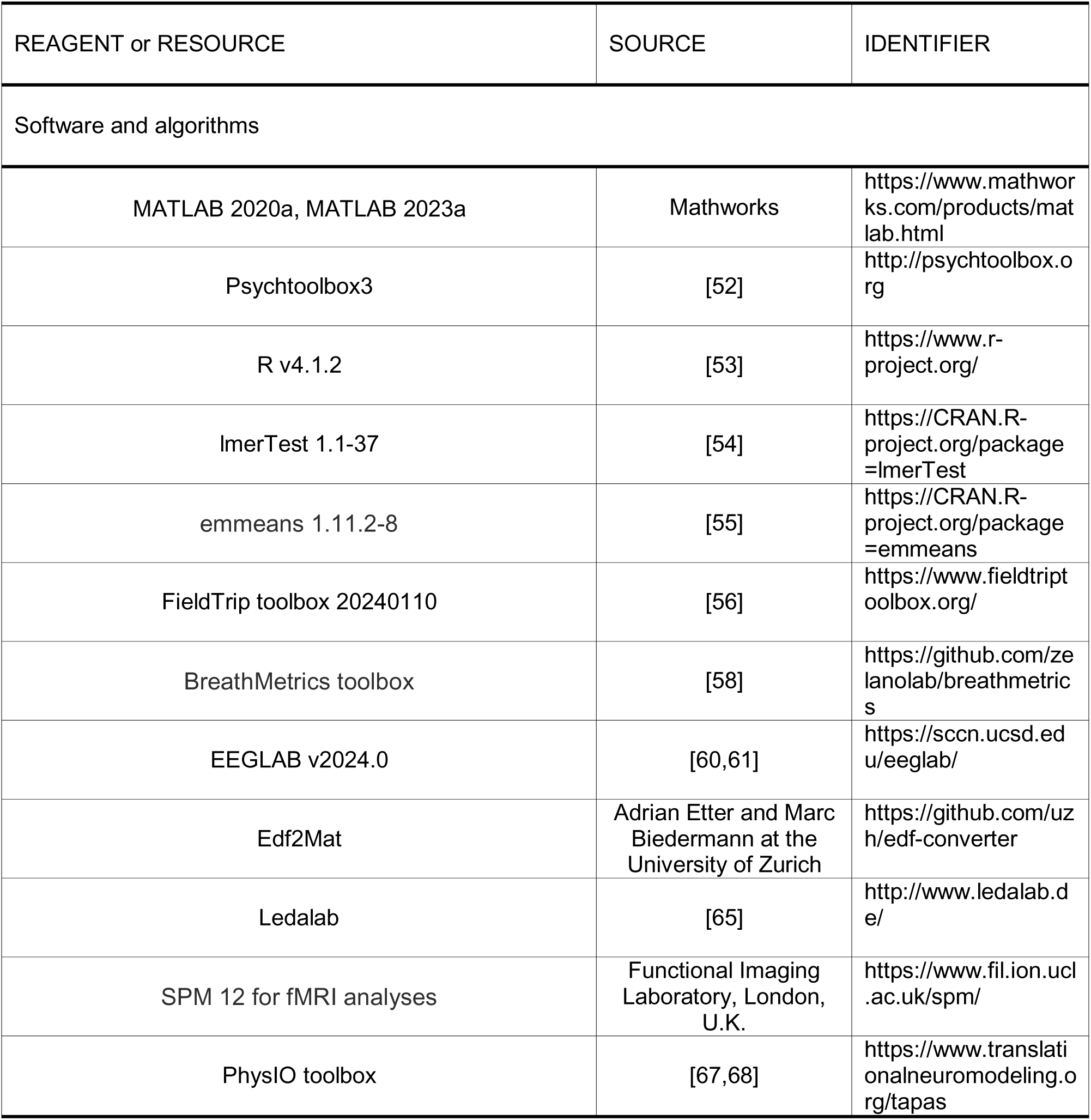

## LEAD CONTACT AND MATERIALS AVAILABILITY

Data supporting the findings of this study will be made available by the Lead Contact upon reasonable request, in accordance with institutional and ethical regulations. This includes anonymised behavioural and physiological data, as well as first-level fMRI contrast images relevant to the main analyses.

## EXPERIMENTAL MODEL AND STUDY PARTICIPANT DETAILS

### PARTICIPANT

Forty-nine healthy adults were recruited from the local university community via flyers and online advertisements. All participants provided written informed consent in accordance with the Declaration of Helsinki, and the study protocol was approved by the Ethics Committee of the University of Potsdam.

Participants were required to be between 18 and 40 years old, fluent in German, right-handed, and to have a body mass index (BMI) between 18 and 25 kg/m². To ensure consistent autonomic and respiratory baselines, individuals were excluded if they reported a history of psychiatric, neurological, cardiovascular, pulmonary, or metabolic disorders; regular medication use; current infection or excessive stress; smoking; or engagement in extreme athletic training. Additional exclusions included abnormal baseline breathing, irregular sleep wake cycles (e.g., night-shift work). MRI eligibility was assessed using standard institutional safety screening.

Eight participants were excluded from all analyses for the following reasons: one participant completed the task using an incorrect breathing pacing bar due to an incorrect assignment of respiratory settings, one was found to have incidental brain abnormalities on structural MRI scanning, one reported visual impairment that prevented accurate perception of task stimuli, one repeatedly fell asleep during the task, and four failed to follow the instructed breathing rhythm during scanning. The final behavioral sample included 41 participants (n=41; 24 female, mean ± SD age = 24.78 ± 4.93 years).

Due to modality-specific signal quality criteria, the number of participants included in each analysis varied. For HRV, electrocardiogram (ECG) was used when available. For participants without usable ECG data, high-quality Photoplethysmography (PPG) recordings were used to derive inter-beat intervals and systolic peaks for RMSSD, RespHRV, and cardiac phase estimation used in fMRI preprocessing. The final HRV sample included 35 participants: 19 with ECG and 16 with PPG. fMRI analyses were limited to these 35 participants, who met both imaging quality and cardiac signal criteria.

## METHOD DETAILS

### Preregistration and deviations

This study was preregistered on the Open Science Framework (https://osf.io/4cbfz). We provide a dedicated supplementary table (Table S13), for each preregistered hypothesis and planned analysis: (i) the preregistered specification, (ii) the corresponding implementation in the current manuscript, (iii) whether there was a deviation, and (iv) the rationale for any deviation.

### Experimental design

#### Breathing manipulation procedure

We used two breathing conditions to manipulate participants’ breathing rhythms: eupnea and prolonged exhalation. Both techniques involved inhaling through the nose and exhaling through pursed lips, with the abdomen rising during inspiration and lowering during expiration. To pace participants’ breathing rhythm, a bar was presented on a screen. The green bar filled when participants were asked to inhale and emptied when they were supposed to exhale. The prolonged exhalation and eupnea conditions differed in their duration and ratio of inspiration and expiration. In the prolonged exhalation condition, participants were instructed to inhale and exhale at a fixed ratio of 2:8 seconds, as previously recommended^51^.

Before the experiment, the individual’s baseline breathing pattern was measured, which was then used for the visual instruction of the eupnea condition. Participants were instructed to breathe normally and calmly (Figure 1A). Specifically, mean inhalation and exhalation durations were computed from 20 valid respiratory cycles following an initial 5-cycle adaptation period, yielding individualized inhale-to-exhale ratios for the eupnea condition. After this calibration, participants practiced both breathing techniques, eupnea and prolonged exhalation, alternating every 3 minutes to learn to synchronize their breathing with the visual cues used during the task.

#### Decision-making task

Participants performed a decision-making task, namely a risky choice paradigm based on a previously established paradigm^11^. Each trial began with a fixation cross (1–9 s), followed by a visual display showing a pair of potential monetary rewards and losses (Figure 1C). Participants had up to 3 seconds to decide whether to accept or reject the presented gamble. Responses were made using a four-point scale: *strongly accept, weakly accept, weakly reject,* or *strongly reject*.

Task presentation and response collection were implemented using Psychtoolbox in MATLAB^52^. During each decision, participants followed a designated breathing rhythm, guided by a visual progress bar on the right side of the screen. They were informed that the probability of winning or losing each gamble was 50%, as in a fair coin toss^9^. Reward amounts ranged from €10 to €30, and loss amounts from €5 to €15. No feedback was provided after each decision, to minimize external interference and maintain consistent task engagement.

Each breathing condition consisted of 120 trials, evenly distributed across three blocks each condition. To ensure incentive-compatible decisions, three trials were randomly selected at the end of the experiment, and participants received or lost 10% of the respective gamble outcomes. Participants could not lose more money than they could gain. Each block lasted approximately 5 minutes. The three blocks of each breathing condition were presented in a continuous sequence, and the order of conditions was counterbalanced across participants (three blocks of prolonged exhalation followed by three of eupnea, or vice versa).

### Behavioral data analysis

#### Behavioral modeling (GLMM)

To examine the effect of breathing condition on decision-making, we modeled the probability of acceptance as a function of breathing condition, gain magnitude, and loss magnitude. We dichotomised responses into a binary variable: accept (strongly or weakly accept) versus reject (strongly or weakly reject).

The decision variable was analysed using glmer with a binomial family and logit link in R (v4.1.2). The model included fixed effects for breathing condition (prolonged exhalation = 1, eupnea = −1), z-standardised reward magnitude, z-standardised loss magnitude, and their interactions with condition. Age, sex, and condition order were entered as covariates.

#### Random-effects structure selection

We conducted a systematic model comparison to determine an appropriate random-effects structure (Table S7–9). Based on AIC/BIC comparisons and established recommendations to balance model complexity and parameter identifiability, we selected model M6, which includes random intercepts and random slopes for condition, reward_z, and loss_z, with correlations. This structure captures all manipulated within-subject main effects while ensuring model convergence and stable parameter estimation.

#### Formal comparison of reward vs. loss modulation

To directly test whether prolonged exhalation differentially modulated gain versus loss sensitivity, we performed a likelihood-ratio test (LRT) comparing the full model against a constrained model forcing the condition × reward_z and condition × loss_z coefficients to be equal.

### Control analyses for alternative explanations

#### Choice entropy

To assess whether prolonged exhalation reduced decision noise, we computed Shannon entropy as a model-free measure of choice consistency. Trials were binned into five quantiles based on z-standardized EV, and entropy was averaged across the central bins for each participant and condition, excluding floor and ceiling effects.

#### Lapse-rate estimation

To further examine potential changes in stimulus-independent responding, we fitted lapse-augmented logistic choice models in which a participant-specific lapse parameter captured random responding independent of value. Lapse rates were estimated separately for each condition at the individual level.

#### Response time analyses

Response times (RTs) were log-transformed and analysed using a linear mixed-effects model including condition × reward_z and condition × loss_z interactions. To test whether RT differences mediated the observed choice effects, trial-wise log-transformed RTs were added as a covariate to the main choice GLMM, and attenuation of the condition × reward_z effect was assessed.

### Physiological data collection and analysis

#### Physiological data acquisition

Physiological signals, including ECG, Electrodermal Activity (EDA), and respiration, were recorded simultaneously using a BrainAmp MR-compatible amplifier (Brain Products GmbH, Germany) and BrainVision Recorder software (version 1.21.0303). All signals were sampled at 5,000 Hz with 16-bit resolution and recorded in DC mode. A three-channel configuration was used: ECG was acquired via a bipolar chest setup; EDA was recorded from the non-dominant hand using a GSR-MR module (resolution: 0.006104 µS); and respiration was measured via a thoracic belt (resolution: 0.1526 arbitrary units). PPG signals were simultaneously recorded at 400 Hz using the Siemens physiological monitoring unit integrated with the MRI scanner. Pupil diameter was recorded at 500 Hz using an MRI-compatible EyeLink 1000 eye-tracker (SR Research Ltd., Canada), positioned at the rear of the scanner bore and tracking the participant’s right eye throughout the task.

#### Respiration

Respiration was recorded using a thoracic belt positioned around the upper abdomen. Then raw signals were low-pass filtered at 1 Hz using a third-order finite impulse response (FIR) filter via the FieldTrip toolbox^56^. The data were downsampled to 100 Hz, detrended, and baseline corrected. Outliers were interpolated using a median-based approach and smoothed with a Savitzky-Golay filter^57^. Inhalation and exhalation durations were extracted cycle-by-cycle using a peak detection algorithm implemented in the BreathMetrics toolbox^58^.

As a quality control measure, each block was evaluated based on the proportion of respiratory cycles with inhalation/exhalation ratios falling within an acceptable range: for the prolonged exhalation condition, a fixed target ratio of 2:8 seconds (inhalation/exhalation = 0.25) was used. The eupnea condition used the individually estimated inhalation to exhalation ratios as a target. In both conditions, a cycle was considered valid if its ratio fell within ±60% of the target ratio.

#### Electrocardiogram (ECG)

ECG was recorded via cutaneous electrodes placed below the clavicle in a bipolar chest configuration. Preprocessing focused on artifact removal and the extraction of inter-beat intervals (IBIs) for heart rate variability (HRV) analysis. Gradient artifacts were corrected using a custom MATLAB script that utilized EEGLAB functions to implement a volume-locked template subtraction procedure^59,60,61^. Scanner volume onset markers were used to extract synchronized ECG segments from each volume, and artifact templates were constructed and subtracted to preserve physiological signals while removing scanner-related noise. Incomplete final volumes were excluded.

Following artifact correction, ECG signals were bandpass filtered between 1 and 100 Hz (fourth-order FIR) to remove baseline drift and high-frequency noise. R-peaks were then identified using a semi-automated MATLAB pipeline that combines z-score normalization, adaptive thresholding, and template matching.

To examine HRV under different breathing conditions, we focused on time-domain measures, specifically the RMSSD and RespHRV, rather than frequency-domain metrics. This decision was based on methodological considerations: time-domain indices are more robust to non-stationary fluctuations and short block-wise recordings and are less confounded by irregular or unbalanced breathing patterns such as those induced by the prolonged exhalation intervention^26,29^.

The final corrected IBI time series were used to compute RMSSD for each breathing condition as a time-domain index of cardiac parasympathetic modulation. In participants where ECG recordings were incomplete or noisy, IBIs were extracted from concurrently recorded PPG data using the same peak detection procedure, and RMSSD was computed accordingly^62,63^.

RespHRV was quantified using a peak-to-trough (P2T) approach based on heart rate fluctuations across the respiratory cycle, following established procedures^5,64^. RespHRV was defined as the difference in heart period between inhalation and exhalation phases and was averaged across trials within each breathing condition.

#### Electrodermal Activity (EDA)

EDA was measured via electrodes placed on the index and middle fingers of the non-dominant (left) hand. The raw signal underwent preprocessing to remove scanner-related artifacts and extract tonic components.Gradient artifacts were removed using the FASTR algorithm as implemented in EEGLAB^60,61^, which adaptively subtracts scanner-periodic noise. The corrected signal was then processed in Ledalab^65^ using a standardized batch pipeline. Specifically, the data were low-pass filtered using a first-order Butterworth filter with a cutoff at 5 Hz, and downsampled from 5,000 Hz to 100 Hz to reduce file size and improve processing efficiency. Adaptive smoothing was applied to further reduce noise. After preprocessing and artifact rejection, EDA data of sufficient quality were retained for 26 participants, who were included in the final analysis.

Continuous Decomposition Analysis was used to separate tonic and phasic components of the EDA signal. Only the tonic component (skin conductance level) was analyzed in the present study as an index of sympathetic arousal. Tonic SCL values were averaged across blocks for each participant and breathing condition.

#### Photoplethysmography (PPG)

Pulse signals were recorded using the Siemens physiological monitoring unit integrated with the MRI scanner, sampled at 400 Hz. The signal was band-pass filtered (0.5–5 Hz, fourth-order Chebyshev Type II) and linearly detrended to remove baseline drift and isolate cardiac-related fluctuations. Systolic peaks were detected using the same semi-automated MATLAB pipeline used for ECG, which combines z-score normalization, adaptive thresholding, and template matching. All detected peaks were visually inspected and manually corrected as needed, including the adjustment or replacement of implausible IBIs.

PPG was used exclusively in participants for whom ECG data were unavailable or of insufficient quality. In these cases, IBIs derived from the PPG signal were used to compute RMSSD, consistent with the ECG analysis.

#### Pupil diameter

Pupil diameter was recorded at 500 Hz using an MRI-compatible EyeLink 1000 eye-tracker (SR Research Ltd., Canada), positioned at the rear of the scanner bore and tracking the participant’s right eye throughout the task. All blocks were conducted under identical lighting conditions, and the visual content (e.g., fixation cross and stimuli) was the same across conditions, ensuring comparable visual input.

Raw pupil data were preprocessed using a custom MATLAB pipeline. The data were converted using the Edf2Mat MATLAB Toolbox designed and developed by Adrian Etter and Marc Biedermann at the University of Zurich. Blinks were identified using EyeLink event markers and periods of zero-valued pupil size. These segments were excluded, and missing values were linearly interpolated to reconstruct a continuous time series. The signal was then downsampled to 100 Hz and corrected for blink-edge artifacts by estimating a peak envelope using spline interpolation with a minimum peak separation of 200 ms. The resulting trace was low-pass filtered using a fourth-order zero-phase Butterworth filter with a cutoff frequency of 8 Hz. Finally, pupil size was baseline-corrected by subtracting the mean value from the first 10 seconds of each block to account for between-subject variability. Pupil data were then visually inspected for quality and retained only if deemed usable based on signal stability and absence of gross artifacts. After preprocessing, pupil data of sufficient quality were retained for 28 participants and included in subsequent analyses.

Average pupil diameter was computed per condition and used as an index of sympathetic arousal.

#### Physiological data analyses

To assess condition differences in physiological signals, we used two-tailed paired t-tests on subject-level condition means. Each subject contributed one average value per condition, enabling a direct within-subject comparison. Effect size was calculated as Cohen’s d for paired samples (mean difference divided by the standard deviation of differences).

#### fMRI data acquisition, preprocessing, and statistical analysis

Whole-brain functional MRI data were collected on a Siemens MAGNETOM Prisma 3T scanner equipped with a 32-channel head coil at the Center for Cognitive Neuroscience Berlin of the Freie Universität Berlin. Functional images were acquired using a T2-weighted echo-planar imaging (EPI) sequence with simultaneous multi-slice acquisition (multi-band factor = 4). The acquisition parameters were as follows: repetition time (TR) = 750 ms, echo time (TE) = 30 ms, flip angle = 65°, 40 axial slices, voxel size = 3 × 3 × 3 mm³, field of view (FOV) = 192 × 192 mm, and phase encoding direction posterior-to-anterior. Multiband acceleration was used to increase temporal resolution. High-resolution anatomical images were obtained using a T1-weighted MP-RAGE sequence (TR = 1900 ms; TE = 2.52 ms; flip angle = 9°; 176 slices; voxel size = 1 × 1 × 1 mm³; FOV = 256 × 256 mm) with GRAPPA acceleration. Blocks were excluded from first-level analysis if head motion exceeded 3 mm translation or 3° rotation on any axis. Based on this criterion, block-level exclusions occurred in 8 participants, of whom 2 had already been excluded due to missing physiological data. The remaining 6 participants were retained in the analysis with partial block-level data (8 blocks total). After combining exclusions from head motion and physiological signal quality, the final fMRI analysis sample comprised 35 participants.

All image preprocessing and analyses were performed using SPM12 (The Wellcome Department of Imaging Neuroscience, Institute of Neurology, London, UK; https://www.fil.ion.ucl.ac.uk/spm/software/). Functional volumes were first slice-time corrected using slice acquisition times provided by the scanner, with the middle slice as the reference. Images were then realigned to the mean image of each run to correct for head motion. Each participant’s T1-weighted anatomical image was co-registered to the mean functional image and segmented into gray matter, white matter, and cerebrospinal fluid using SPM’s unified segmentation procedure. Spatial normalization to the Montreal Neurological Institute standard space was performed using the deformation fields generated during segmentation. Functional images were resampled to 3 × 3 × 3 mm³ voxel size and spatially smoothed with an 8 mm full width at half-maximum Gaussian kernel.

To rule out the effect of general physiological signals, we performed physiological noise correction using RETROICOR^66^ based on concurrently recorded cardiac and respiratory signals. Physiological regressors were generated using the PhysIO toolbox^67,68^, modeling cardiac phase (third-order Fourier expansion), respiratory phase (fourth order), and their first-order interaction. This resulted in 18 physiological regressors per block. In addition, the six motion parameters estimated during realignment (three translations and three rotations) were included as nuisance covariates in the first-level GLM to account for non-neural sources of BOLD signal variability and remove physiological noise unrelated to neural activity.

To investigate neural responses to rewards and losses, we specified a GLM in which the onset times of options were modeled as events and parametrically modulated by two regressors: the magnitude of rewards and the magnitude of losses. All regressors were convolved with the canonical hemodynamic response function. First-level contrast images were computed to capture condition-specific neural sensitivity to reward and loss magnitude. Reward and loss were entered as separate parametric modulators of the option-onset regressors without orthogonalisation, so that each regressor captures variance uniquely associated with its respective magnitude.

At the group level, contrast images were entered into a second-level random-effects analysis. To assess whether individual differences in cardiac parasympathetic regulation predicted changes in neural sensitivity to prospective reward, we conducted a second-level whole-brain regression analysis. The input for this analysis was the first-level contrast images representing the parametric reward effect difference between prolonged exhalation and eupnea. For each participant, the within-subject difference in heart rate variability (ΔRMSSD (prolonged exhalation - eupnea)) was computed and entered as a continuous covariate. Whole-brain regression was performed using a voxel-level threshold of p < 0.001 (uncorrected), followed by cluster-level family-wise error (FWE) correction at p < 0.05.

## ACKNOWLEDGMENTS

This study was funded by German Center for Mental Health (Deutsches Zentrum für Psychische Gesundheit BMBF 01EE2301E), German Center for Diabetes Research (Deutsches Zentrum für Diabetesforschung 82DZD03D03); and BMBF DecEnt 01GP2210C, all acquired by SQP. IR was supported by the Marie Skłodowska-Curie Action (MSCA) BRAINSTOM (grant agreement No 101028203)

## AUTHOR CONTRIBUTIONS

SQP, SR, WH conceptualized the study, WH, MS, LYL acquired data, SQP, WH, GB, MS, IR, MP, FM, BK, analysed the data, SQP, WH, SR, GB, MS, IR, MP, FM, BK, PNCM wrote, reviewed, and edited the manuscript.

## DECLARATION OF GENERATIVE AI AND AI-ASSISTED TECHNOLOGIES

During the preparation of this work, the authors used ChatGPT-4o in order to improve clarity, refine language, and check grammar. After using this tool or service, the authors reviewed and edited the content as needed and take full responsibility for the content of the publication.

## SUPPLEMENTAL INFORMATION

Document Supplemental Information.

